# Jumbo phages possess independent synthesis and utilization systems of NAD^+^

**DOI:** 10.1101/2024.04.18.590177

**Authors:** Cunyuan Li, Kaiping Liu, Chengxiang Gu, Ming Li, Ping Zhou, Linxing Chen, Shize Sun, Xiaoyue Li, Limin Wang, Wei Ni, Meng Li, Shengwei Hu

## Abstract

Jumbo phages, phages with genomes >200 kbp, contain some unique genes for successful reproduction in their bacterial hosts. Due to complex and massive genomes analogous to those of small-celled bacteria, how do jumbo phages complete their life cycle remain largely undefined. In this study, we assembled 668 high-quality jumbo phage genomes from over 15 TB of intestinal metagenomic data from 955 samples of five animal species (cow, sheep, pig, horse, and deer). Within them, we obtained a complete genome of 716 kbp in length, which is the largest phage genome so far reported in the gut environments. Interestingly, 174 out of the 668 jumbo phages were found to encode all genes required for synthesis of NAD^+^ by the salvage pathway or Preiss-Handler pathway, referred as NAD-jumbo phage. Besides synthesis genes of NAD^+^, these NAD-jumbo phages also encode at least 15 types of NAD^+^-consuming enzyme genes involved in DNA replication, DNA repair, and counterdefense, suggesting that these phages not only have the capacity to synthesize NAD^+^ but also redirect NAD^+^ metabolism towards phage propagation need in hosts. Phylogenetic analysis and environmental survey indicated NAD-jumbo phages are widely present in the Earth’s ecosystems, including the human gut, lakes, salt ponds, mine tailings, and seawater. In summary, this study expands our understanding of the diversity and survival strategies of phages, and in-depth study of the NAD-jumbo phages is crucial for understanding their role in ecological regulation.

## Introduction

Bacteriophages (phages), as viruses that infect bacteria, are key components in natural microbiomes because they can shape microbial composition and structure, alter host’s metabolisms, spread antibiotic resistance genes, and mediate gene transfer [1–3]. Metagenomic investigations have showed that diverse and abundant gut virome was observed in livestock gut [4, 5]. Recent studies also demonstrated that phages in livestock microbiome may impact the major functions of gut such as feed digestion, microbial protein synthesis, methane emissions [6, 7]. However, we still lack a deep understanding of how microbes interact with phages to affect gut ecosystem.

Unlike the small genome size commonly understood for phages, some phages have enormous genomes exceeding 200 kbp, referred to as jumbo phages [8], and the largest phage genome reported to date reaches 735 kbp [9]. These jumbo phages are distributed in various environments [9, 10]. Current research indicates that jumbo phages encode auxiliary metabolic genes, such as genes for methane monooxygenases (*pmoC*), to improve the host’s survival ability [11]. Jumbo phages may redirect biosynthesis to phage-encoded functions by CRISPR-induced interception of the earliest steps of hosts translation [9]. Some megaphages with genome > 500 kbp infect *Prevotella* species enriched in the gut microbiomes, may be common but largely overlooked components of human and animal gut microbiomes [12, 13]. However, due to the difficulty in assembling large genomes, the number of reported jumbo phages is still relatively scarce in animal guts [14]. We still know very little about the general characteristics of jumbo phages in animal microbiomes.

In this study, we aim to investigate the diversity, genetic repertoires, and survival strategies of jumbo phages in animal microbiomes. We conducted large-scale data mining for identifying jumbo phages genomes by analyzing over 15 TB of metagenomic data from 955 intestinal samples of five animals (cow, sheep, pig, horse, and deer). We identified 174 jumbo phages with all genes required for synthesis of NAD^+^, referred as NAD-jumbo phage. These NAD-jumbo phages were further characterized by unique genetic repertoires including NAD^+^-dependent enzymes, DNA replication machinery, DNA repair-related enzymes, and counterdefense systems. We also investigated the distribution of NAD-jumbo phages in natural environments and human microbiome, and detected the appearance of NAD-jumbo phages across Earth’s ecosystems. Our study provides new insights into the genomic characteristics and survival strategies of jumbo phage, and novel path forward to exploring bacteria-phage interactions.

## Results

### Basic features of jumbo phages identified in animal gut microbiomes

To characterize the diversity of jumbo phages in animal gut, we analyzed 955 metagenomes from 15.24 TB dataset including 8.71 TB for cattle, 1.96 TB for horses, 1.25 TB for pigs, 2.75 TB for sheep and 0.56TB for deer (Supplementary Table 1). Based on previous reports [9, 15], we have developed a rigorous phage detection pipeline to identify phages from the metagenomic datasets (Figure S1, see Methods). A total of 1001 *de novo* assembled contigs longer than 200 kbp were classified as jumbo phages (Supplementary Table 2). To improve the quality of subsequent analysis, 668 non-redundant jumbo phages with high-quality (completeness >90%, 378 in total) and complete (290 in total) genomes were selected to form a high-quality gut jumbo phages set of animals.

The 668 jumbo phage genomes were distributed in five animal species (Figure S2 and S3), representing as the largest number of intestinal jumbo phage genomes reported so far to our knowledge. There are 10 phages whose genome size was >500 kbp, which can be classified as megaphage. These megaphages were found in the intestines of cattle, sheep, horses, and pigs, suggesting that the animal intestine may be a megaphage reservoir (Figure S2). Two bioinformatically validated circular and complete phage genomes, ERR20279001_716809 (Houyi_20) and SRR44355611_713556 (Houyi_34), are 716.8 kbp and 713.6 kbp in length, respectively, and are the largest phages reported to date in the gut (Figure S4). The previously reported largest intestinal phage genome is 660 kbp in length from horse gut samples [13]. Both phages with genome length of 716.8 kbp and 713.6 kbp we are derived from the rumen metagenome data of cattle, but the average nucleotide homology (ANI) of the two phage genomes is 93.93%, indicating that the two phages are different species, which represents the diversity of megaphages in the rumen. The median genome size of the 668 high-quality gut jumbo phages of animal was 263.28 kbp, which were much bigger than those of IMG/VR and previous reports [16, 17]. In total, the genome set here expand the catalogue of jumbo phage and provide a rich source for jumbo phage biology.

In addition, the jumbo phage genomes averagely encode 23 tRNA genes with predicted functions in translation (Supplementary Table 3), and 82.29% of them with at least 5 tRNA genes. SRR12268561_1_289927 (Houyi_39) encodes up to 68 tRNAs, which cover the full range of amino acids required for protein synthesis. Interestingly, a suppressor tRNA called sup-tRNA was encoded by some jumbo phage genomes, which can induce translational readthrough of nonsense mutations. It seems that they may help increase the translation efficiency of phage genes under stress [16]. The tRNA sequence encoded by the jumbo phage is different from that of the host (Figure S5), which may help phages redirect host’s ribosomes to make more of their own proteins [9].

### Diversity and hosts

As jumbo phage had expanded genomes sizes, the evolutionary relatedness of our newly assembled and published jumbo phages was unclear. Since the commonly used classification marker genes for phages cannot cover all groups, preliminary grouping can usually be performed according to gene content [19]. A network including the 668 jumbo phages and published phages was generated to group the newly-assembled jumbo phages, which showed their novelty (Figure S6). To further investigate their evolutionary relationship, we constructed phylogenetic trees using major capsid protein (MCP) and terminase large subunit (TerL) (Figure 1). Out of the 668 jumbo phages, 143 have both MCP and TerL, and 291 have at least one of them (Supplementary Table 4-5). The proteins of the jumbo phages reported here were well clustered with high guidance support, thus defining the evolutionary clades. Most of jumbo phages fall into one of 5 clades in MCP and TerL trees. We named these 5 clades of jumbo phages as Kuafu_Jumbo phage, Houyi_jumbo phages, Jingwei_jumbo phages, Pangu_jumbo phages and Zhurong_jumbo phages (according to the characters in traditional Chinese mythology) (Figure 1, Supplementary Table 5). Within each clade, phages were sampled from different animal species from different countries in the world, suggesting the diversification of these jumbo phages across gut environments (Figure 1). The megaphages (genome size > 500 kbp) from different animal intestines clustered into the same clade, suggesting that megaphages in animal intestines may share their ancestor (Figure 1). In addition, no typical virus MCP sequences were found in Zhurong_jumbo phage clade.

**Figure 1.**
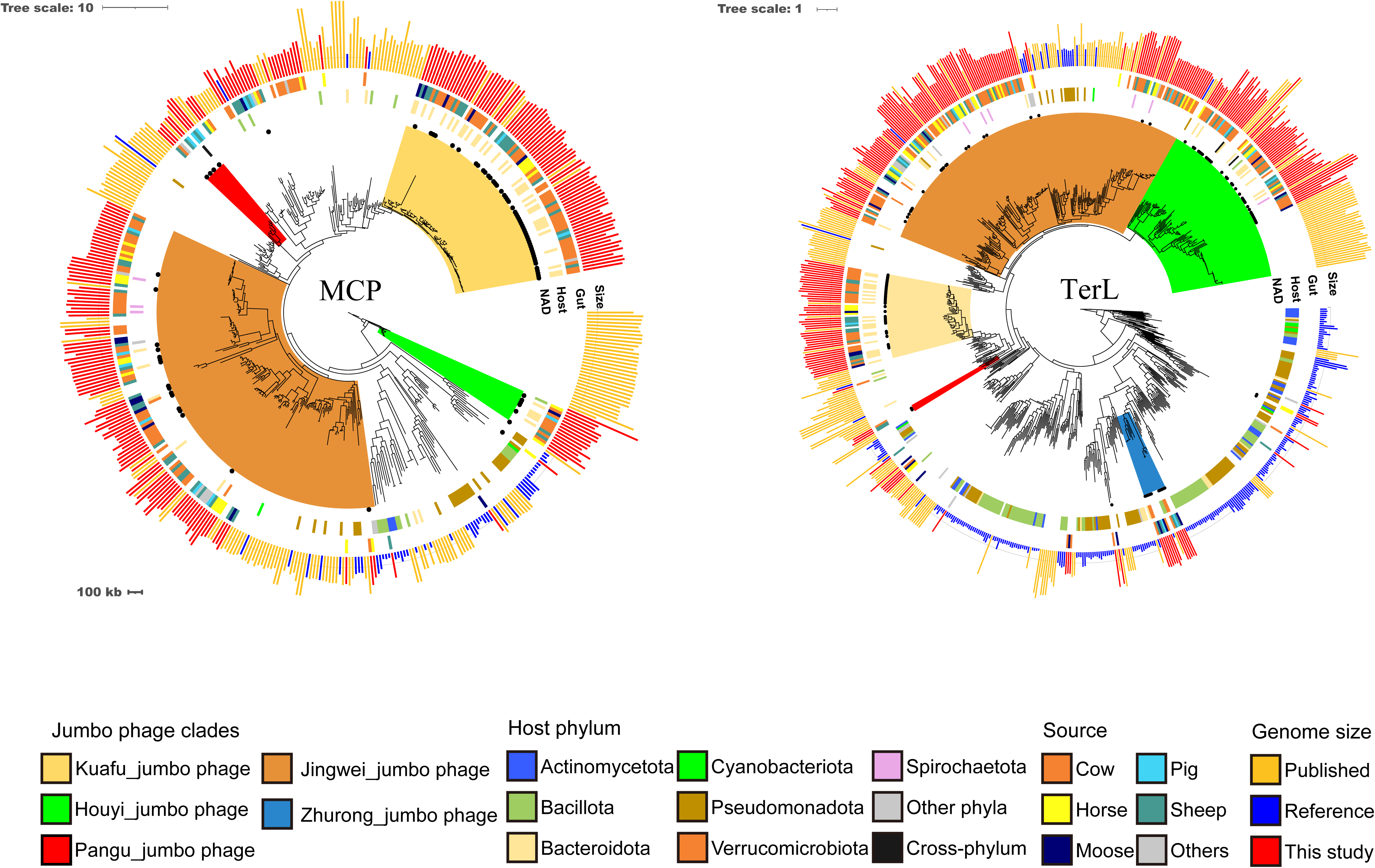
Phylogenetic tree of jumbo phages. Phylogenetic tree was constructed by major capsid proteins (MCP, left) and large terminase (TerL, right) sequences identified in this study. Homologs references were collected from published jumbo phages [9] and VOG database. From inner to outer ring, the concentric circles indicate the presence of NAD-jumbo phage in this study, host phylum, gut source, and genome size. Jumbo phage clades indicate five different clades of phages identified in this study. Host phylum indicates the predicted host phylum of jumbo phages using CRISPR spacer analysis. Gut source represents which animal guts sampled to obtain jumbo phages in this study. Genome size is sequence length of phage genomes used in the tree. Different colors represent different types of clades, host phylum, gut source, and genome size in the legends.

Predicting the cellular host of the viruses is important for understanding the virus-host interactions [20]. We used CRISPR spacer targeting to detect phylum-level link between host and the jumbo phages by collecting metagenome assembly genomes (MAGs) assembled from the animal gut microbiomes, then constructed an animal gut MAG dataset containing 29,599 bacterial and archaeal MAGs (Supplementary dataset 1). We extracted 265,818 CRISPR spacers (Supplementary Table 6) from the datasets, which were rigorously matched to the assembled jumbo phage genomes. We detected 127 links between 123 newly assembled jumbo phages and 10 bacteria phyla. Bacteroidota (63) and Bacillota (34) were the most common host phyla of jumbo phages (Figure 1, Supplementary Table 6).

### Jumbo phages possess independent synthesis pathways of NAD^+^

Annotation of the new jumbo phage genomes revealed they contains a large number of genes involved in DNA replication, transcription, DNA modification, energy synthesis and metabolism (Supplementary Table 4). We are attracted to the NAD^+^ synthesis pathway related genes. The KEGG and gene blast analysis revealed that 174 jumbo phages have all the genes necessary for biosynthesis of NAD^+^ by salvage pathways or Preiss handler pathway (Figure 2A, Supplementary Table 7). The 174 jumbo phages with at least one complete pathway of NAD^+^ synthesis were named as NAD-jumbo phages.

**Figure 2.**
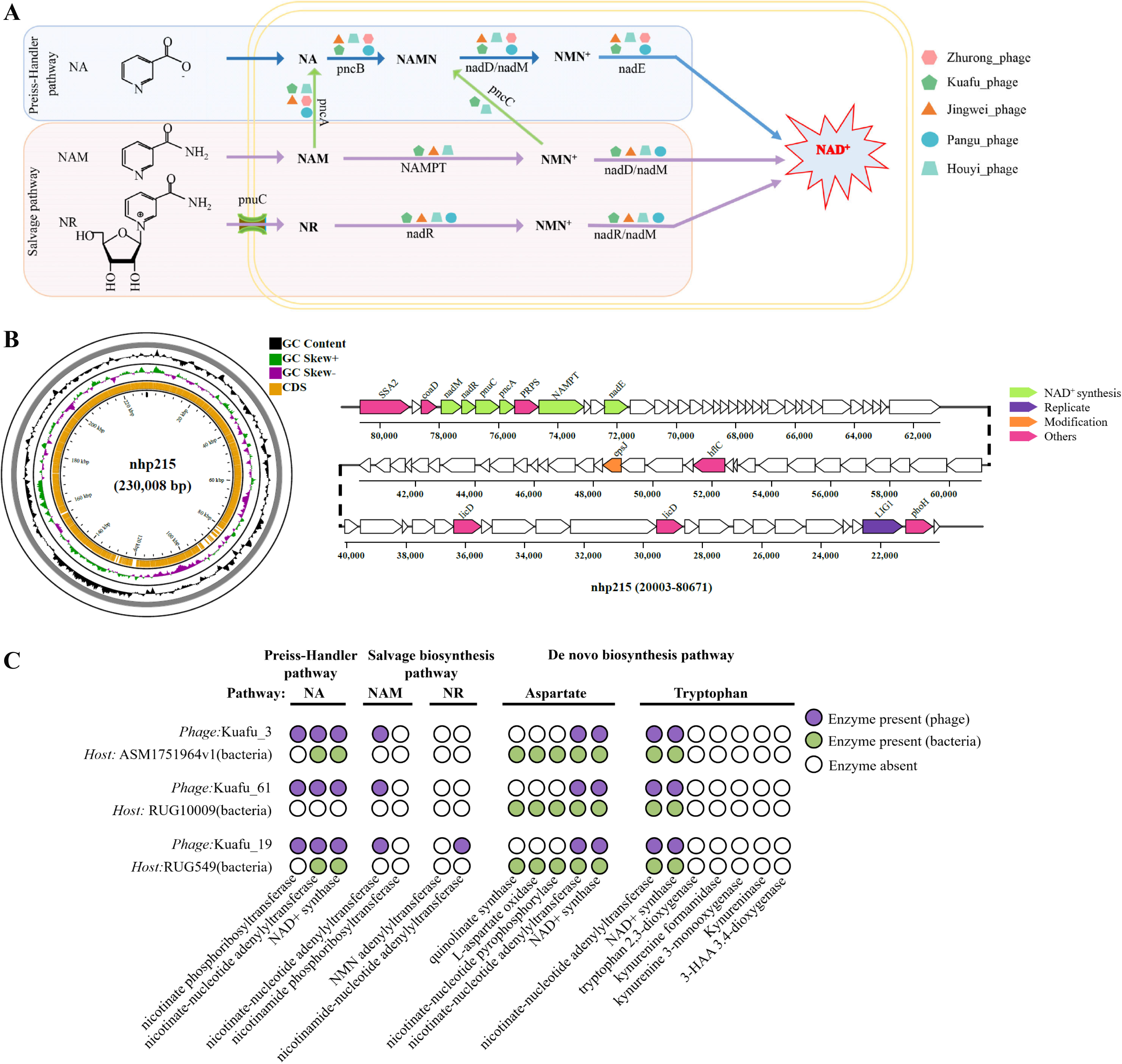
Jumbo phages possess independent synthetic pathways of NAD^+^. **a**, Jumbo phages encode key enzyme genes in each step of NAD^+^ biosynthesis by salvage pathways or Preiss handler pathway. Different colours represent different types of jumbo phages that contain all enzymes necessary for the pathway. The metabolic pathways were obtained from KEGG database. **b,** Genetic maps of NAD-jumbo phage nhp215. Genome circos map (left) and gene content (right) of nhp215. The green graph represents the gene for NAD synthesis. The white graph represents the unannotated gene. **c,** NAD^+^ biosynthesis pathway are complementary between NAD-jumbo phages and their hosts. NAD-jumbo phages are paired with their host bacteria, and their names are displayed on the left. Enzymes involved in each pathway are listed below the column. Green circles and purple circles represent the presence of enzymes encoded by phages and their hosts, respectively.

Interestingly, our 10 largest phages (536-716 kbp in length) all NAD-jumbo phages. Moreover, the phages with genome of 716 kbp, 713 kbp, 696 kbp and 648 kbp contained two different types of NAD^+^ synthesis pathways, such as containing both NAM salvage pathway and NR salvage pathway (Supplementary Table 7). We further analyzed whether NAD synthesis pathway is also present in the previously reported the largest complete phage genomes with 634 kbp, 636 kbp, 642 kbp and 735 kbp in length [9]. We found all but the 735 kbp phages have the complete NR salvage pathway, thus are to NAD-jumbo phages as well. However, the 735 kbp phage carry one nicotinate nucleotide adenylyltransferase (nadD) which play a vital rate-limiting step in the NAD^+^ biosynthesis. Previous experiments documented that the overexpression of bacterial nadD improves NAD^+^ synthesis in a variety of bacteria [21–22].

Notably, NAD^+^ synthesis related genes are physically clustered together on phage genomes to form biosynthetic gene clusters (BGC) that recurred in some NAD-jumbo phages (Figure 2B, Figure S7). For example, NAD-jumbo phages nhp215 carry a BGC clustering nadR and nadM genes which constitute complete NR salvage pathway (Figure 2B); Kuafu_71 carry a BGC clustering Pncb, nadD and nadE genes which are all the genes necessary for biosynthesis of NAD^+^ by Preiss handler pathway (Figure 2A, Figure S7). The BGCs provide complete pathway of NAD synthesis, which may allow jumbo phages to synergistically express genes and potentially promote the production of NAD^+^.

Many NAD-jumbo phages encode genes with predicted functions in key regulator of NAD^+^ biosynthesis, especially nicotinamide hosphoribosyltransferase (NAMPT), and nicotinamide riboside transporter (PnuC). NAMPT is the rate-limiting enzyme of NAD^+^ salvage pathways. The 71 out of 174 NAD-jumbo phages encoded NAMPT genes (Table S4), and phage-NAMPT likely accelerate catalysis of NAM to NMN that were used as a precursor for NAD^+^ biosynthesis in hosts (Figure 2A). For PnuC, it is a transporter catalyzing cellular uptake of the NAD^+^ precursor nicotinamide riboside (NR) across bacterial cell membrane. PnuC genes were encoded by 68 out of 174 NAD-jumbo phages (Table S4). PnuC form a biosynthetic gene cluster with nadR and nadM in NAD-jumbo phages nhp215 (Figure 2B), and may transport more NR substrate from outside of the bacterial membrane to phage NAD^+^ synthesis.

We found the NAD^+^ synthesis pathway of some jumbo phages was complementary to that of the bacterial host (Figure 2C). For example, Kuafu_19 has the Preiss handler pathway to use the substrate NA to synthesize NAD^+^, while its host has the *de novo* synthesis pathway to use aspartate as substrate to synthesize NAD^+^ (Figure 2C). The complementarity of NAD^+^ synthesis between some jumbo phages and their hosts has the opportunity to promote the synthesis efficiency of NAD^+^ and improve the concentration of NAD^+^ available to phages or host, which may further facilitate phage propagation.

NAD-jumbo phages were observed to cluster in phylogenetic analysis, which can be roughly divided into 5 categories (Figure 1). These 174 NAD-jumbo phages were mainly clustered in the clade of Kuafu_jumbo phage and Houyi_jumbo phage. In particular, Kuafu_jumbo phage clade is mainly composed of our newly assembled NAD-jumbo phages. Surprisingly, all predicted hosts for Kuafu_jumbo phages were Bacteroidota, suggesting that Phylum-level host specificity of these phages. For Houyi_jumbo phage, the NAD-jumbo phages are the most important part in Houyi_clade. Meanwhile, the largest gut phages ERR20279001_716809 (716 kbp) and SRR44355611_713556 (713 kbp) belong to Houyi_jumbo phage.

### NAD-jumbo phage encodes NAD-dependent enzymes for various biological processes

Considering the crucial role of NAD^+^ in various biological processes including DNA replication, DNA repairs and redox reactions for all organisms [23], NAD^+^ may be highly significant for life cycle of NAD-jumbo phages. We performed genome-wide functional annotation of NAD-jumbo phages. Our NAD-jumbo phage encodes 15 types of NAD^+^-consuming enzymes (in total of 222 NAD-dependent enzymes or NAD^+^ cofactors) (Supplementary Table 8), which may be involved in DNA replication, DNA repair, and NAD^+^-mediated counterdefense, suggesting that NAD-jumbo phages may mediate various biosynthetic process by expressing own NAD-dependent enzymes (Figure 3).

**Figure 3.**
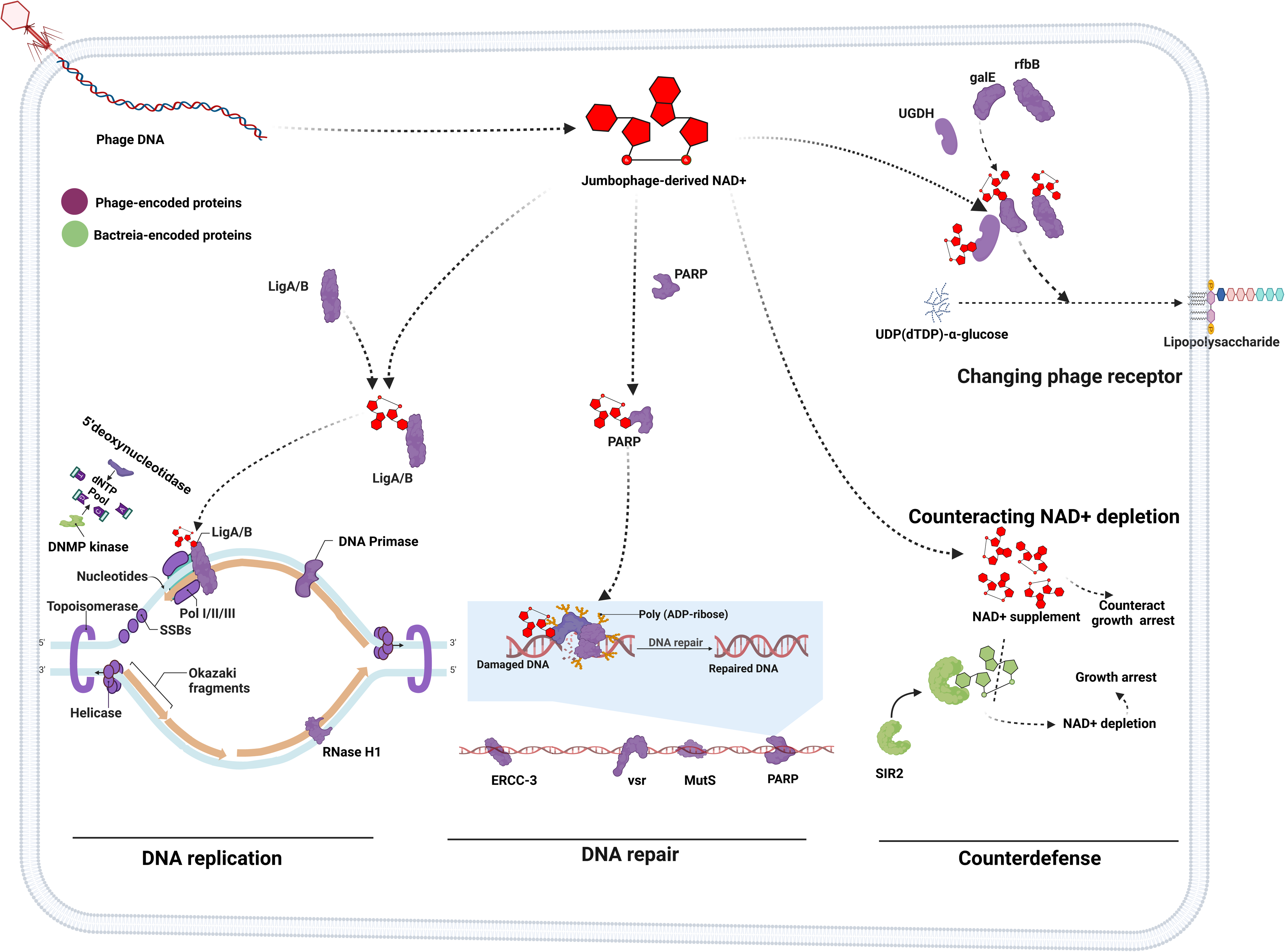
A model for NAD^+^-mediated jumbo phages direction of DNA replication, DNA repair and counterdefense in bacterial hosts. NAD-jumbo phages independently synthesize NAD^+^ and then utilize NAD-dependent enzymes to potentially direct biosynthesis towards phage propagation need in host bacteria. Phage-encoded NAD-dependent enzymes were involved in DNA replication, DNA repair, and counterdefense (NAD^+^ in red, phage-encoded proteins in purple, bacteria-encoded proteins in green). Specially, many NAD-jumbo phages encode almost all genes required for DNA replication and some key genes for initiating DNA repair. LigA/B: NAD-dependent DNA ligase. SSBs, single strand DNA-binding protein; PARP, poly [ADP-ribose] polymerase; ERCC-3, DNA excision repair protein ERCC-3; Vsr, DNA mismatch endonuclease; MutS, DNA mismatch repair protein MutS. Figures generated with BioRender (https://biorender.com/).

It is noted that over half (90/174) of the NAD-jumbo phages contain NAD-dependent DNA ligase (Figure S8, Supplementary Table 7), which is an enzyme necessary for DNA replication and repair. More importantly, these NAD-jumbo phages also encode almost all replication-related genes including DNA topoisomerase, DNA helicase, DNA polymerases family, DNA primase RNase H1, and NAD-dependent DNA ligase (Figure 3), which are key for DNA replication initiation and elongation in prokaryotes [24, 25], and phage may use these proteins to shift DNA biosynthesis towards phage own replication. Therefore, besides potentially independent synthesis of NAD^+^, NAD-jumbo phages may express own NAD-dependent enzymes and replication machinery to direct phage self-replication in bacterial hosts.

Similarly, DNA repair may also be directed towards phage own repair need in bacterial hosts, given that many NAD-jumbo phages carrying genes including DNA damage sensor PARP1 (NAD-dependent type), mismatch recognition factor MutS, as well as excision repair enzymes Vsr and ERCC3 (Supplementary Table 7 and Table 8). Both MutS and NAD-dependent PARP1 could initiate various forms of DNA repair [26, 27]. The Vsr endonuclease and ERCC3 function in nucleotide excision repair, and these two enzymes were likely used for cutting mismatched DNA in phage genomes [28, 29]. NAD-dependent DNA ligase of phage seal nicks in genomes in the final stages of DNA-repair pathways. Together, NAD-jumbo phages may use their own NAD^+^ products, NAD-dependent enzymes and key repair-related factors to direct self-repair in host.

In addition, it is important to note that many NAD-jumbo phages encode NAD-dependent enzyme genes, such as UDP-glucose 6-dehydrogenase (UGDH), UDP-glucose 4-epimerase (galE), and dTDP-glucose 4,6-dehydratase (rfbB), involved in lipopolysaccharide (LPS) biosynthesis (Figure 3, Table S9). Bacteria LPS is a rich and diverse cell wall polysaccharide, which was adopted by phages as receptor for infection [30, 31]. Phage-encoded synthesis of LPS may be able to change the modification of cell wall to prevent the infection of competing phages [32, 33]. The survival adaptability of NAD-jumbo phages may be improved by modifying the host cell wall.

Another function of NAD^+^ derived from jumbo phages may be to counteract bacterial abortive infection defense mediated by NAD^+^ depletion. NAD^+^ depletion triggered by the Sir2 domain cause bacteria suicide to limit phage propagation, which has been well documented in anti-phage defense system [34, 35]. NAD-jumbo phage harbors a diverse array of genes involved in the NAD^+^ salvage synthesis pathway, which are absent in its host (Figure 2C), implying that NAD^+^ may be autonomously synthesized by NAD-jumbo phage. NAD^+^ from NAD-jumbo phage may be used to counteract the host NAD^+^ depletion and maintain the normal level of NAD^+^ for propagation of both host and phage. This suggests that the ability to synthesize NAD^+^ in jumbo phage may be a counter-defense strategy against host NAD^+^ depletion, although this needs further experimental verification.

### NAD-jumbo phages are ubiquitous and diverse across Earth’s ecosystems

Our NAD-jumbo phages spanned five animal species from different countries in the world, reminding us diverse ecosystem distribution of NAD-jumbo phages. To end this, we further analyzed whether NAD^+^ synthesis gene pathways were present in human-associated and environments-associated jumbo phages. Of 863 jumbo phages from human and nature environments microbiome, we identified 181 jumbo phages with complete pathway of NAD^+^ synthesis, which belong to NAD-jumbo phages. We identified distribution of these NAD-jumbo phages across Earth’s ecosystems, including cultivated phages, in human-associated samples (gut and mouth), and in environment-associated samples (freshwater lakes and rivers, marine ecosystems, sediments, hot springs, soil, deep subsurface habitats and built environment) (Figure 4A, Supplementary Table 9 and 11). The NAD-jumbo phages have been isolated and cultivated on different bacteria hosts, indicating that the identification of such NAD^+^ synthesis pathway in our study is not an artefacts of metagenome analyses.

**Figure 4.**
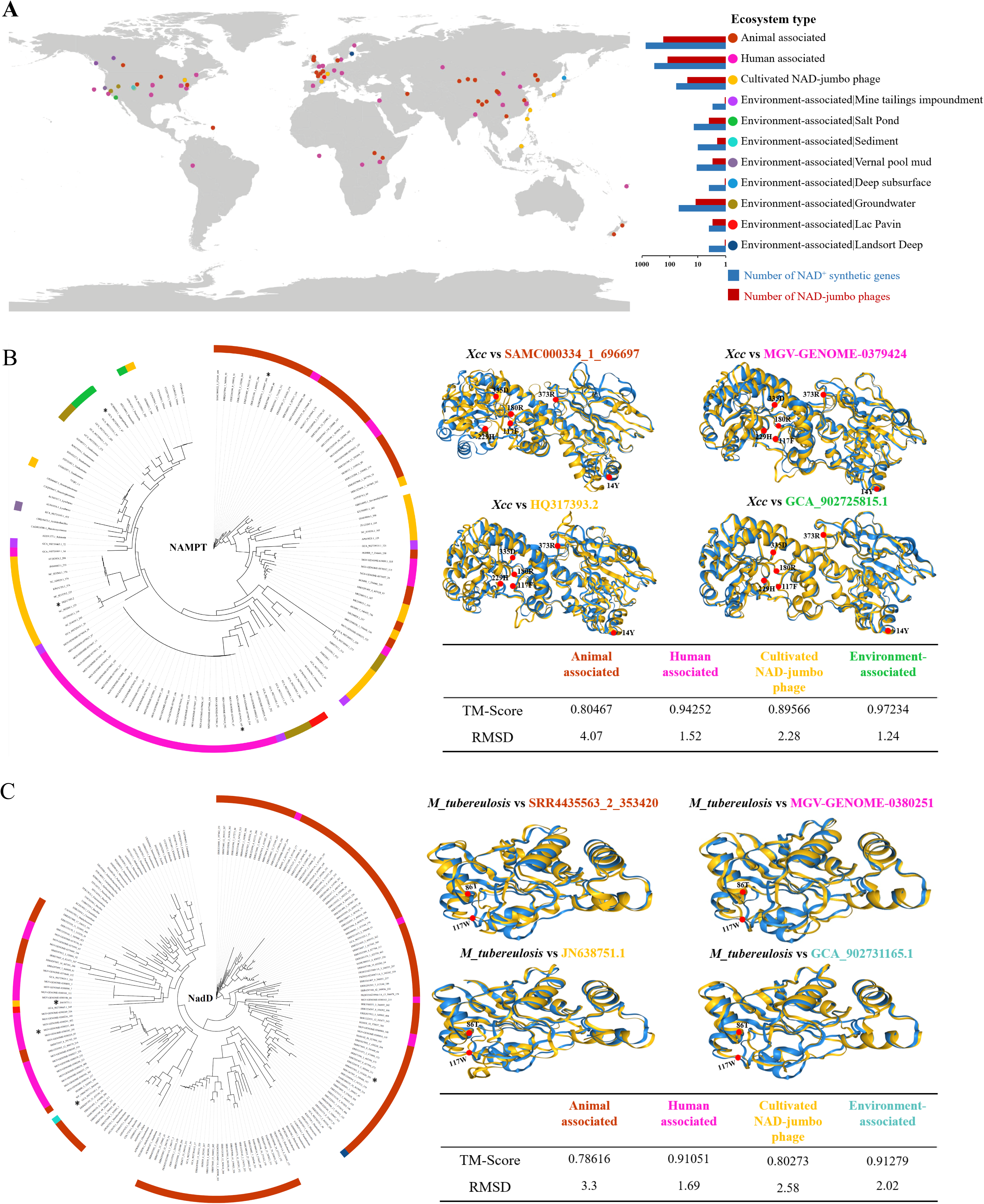
The distubrition of NAD-jumbo phages across Earth’s ecosystems. **a**, Global distribution of NAD-jumbo phages identified in 12 different ecosystems types. The different symbols and colored dots represent different ecosystem types in the legend. **b, c,** Phylogenetic trees (left) and structures (right) of NAD-jumbo phages NAMPT and nadD proteins. Phage-encoded protein sequences were extracted from animal-, human– and environment-associated NAD-jumbo phage genomes. Microbial reference sequences were collected from RefSeq database. Different colors represent ecosystem origin of protein. Asterisk indicates that the sequence is used for protein structure prediction. Bootstrap values greater than 50 are shown on the branches. For protein structures, blue represents reference of bacteria M_tuberculosis NAMPT and Xanthomonas campestris pv. campestris (Xcc) nadD protein structures from cryo-electron microscope results; Yellow represents RoseTTAFold-predicted viral protein structures of NAMPT and nadD from different ecosystem types and cultivated phages. The red dot on protein structures indicates bioactive amino acid residues.

Similar to animal-associated NAD-jumbo phages, the environment– and human-associated NAD-jumbo phages also include NR salvage pathway, NAM salvage pathway, or Preiss handler pathway, showing diversity of NAD^+^ synthesis pathways across ecosystems (Supplementary Table 9). We further characterized the diversity of NAD^+^ synthesis genes by conducting phylogenetic analysis of two NAD^+^ synthesis rate-limiting enzymes (NAMPT and nadD) sequences. In most cases, phylogeny of the NAD^+^ synthesis enzymes from the same ecosystem type display close clustering, supporting previous finding that phage metabolic enzymes are correlate with environment temperature and nutrition for enhancing phage environmental adaptation [36]. In few cases, NAD^+^ synthesis enzymes phylogeny of human-associated phages show close clustering with those of animal associated-phages, and likely explanation is that horizontal gene transfer occurs in NAD-jumbo phages of between animals and human. Although NAD^+^ synthesis genes were diverse across cosystems, RoseTTAFold predicted protein structures and conserved amino acid residues of NAMPT and nadD were consistent with those of bacteria determined by cryo-electron microscope (Figure 4B, C).

Besides the synthesis genes of NAD^+^, the environment– and human-associated NAD-jumbo phages also carried NAD-consuming enzyme genes involved in DNA replication, DNA repair and counterdefense (Supplementary Table S11), which was consistent with those found in the animal gut. For NAD-jumbo phages in human gut, the hosts of these phages are mainly concentrated in Bacteroidota (Supplementary Table 12). NAD-jumbo phages in human microbiomes have complement NAD^+^ synthesis pathway with their hosts (Figure S9). Collectively, phage-encoded NAD^+^ synthesis and utilization systems are widely distributed across Earth’s ecosystems, and potentially assist jumbo phage functions in various environment.

## Discussion

Our study indicates that NAD-jumbo phages are highly abundant and widely distributed across Earth’s ecosystems. We have successfully assembled a total of 1001 jumbo phage genomes. Among of them, 668 non-redundant jumbo phage genomes were high-quality or complete, which are the highest quality and largest quantity of jumbo phage genome resources to date. By integrating 863 jumbo phage genomes from public database and previous reports [9], we form global Jumbo phage database (JumbophageDB), which contains 1864 viral draft genomes across ecosystems (Supplementary dataset 2). This resource further expands the unexplored viral diversity not found in other databases, and it will improve detection of viral reads in microbiome and may help to investigate their roles in gut health and natural environment.

It has been reported that bacteriophage Vibrio KVP40 encoded NAMPT have shown the highest activity, resulting in a threefold increase in production of NAD^+^ precursor (NMN), when compared with eight NAMPTs from different sources of eukaryotes and prokaryotes [37]. NMN, as an effective anti-aging product to increase the NAD^+^ levels, ameliorates the detrimental effects of NAD^+^ reduction with age, and significantly improves longevity as well as to ameliorate age-related complications [38]. KVP40-encoded NAMPT have been used for large-scale production of NMN in engineering bacteria. NAD-jumbo phages include 71 NAMPTs and other hundreds of NMN synthesis genes (nadR, nadM, nadD) (Table S4), which provides a valuable gene repository for producing active NMN by genetically engineered bacteria or yeast.

NAD-jumbo phages possess expanded genetic repertoires including all enzyme genes required for synthesis of NAD^+^ through the salvage pathway and the Preiss-Handler pathway rather than the de novo pathway. It seems to be related to the high efficiency of NAD^+^ synthesis by the salvage pathway and the Preiss-Handler pathway [39]. Two studies have reported that phage-encoded enzymes of NAD^+^ synthesis show highly biological activity in vitro and in bacteria [37, 40], implying that NAD-jumbo phages may synthesize true NAD^+^ in hosts. In addition, NAD-jumbo phages also encode at least 15 types of NAD^+^-consuming enzyme genes involved in DNA replication, DNA repair, and counterdefense, suggesting that phages have the capacity to redirect NAD^+^ metabolism towards phage propagation need in bacterial hosts. The remodeling of NAD^+^ metabolism is aligned with findings for viruses infecting archaea and cyanobacteria, which redirect the chemical energy (ATP) synthesis to increase viral production [41]. The remodeling metabolism of NAD-jumbo phages may represent a survival strategy or counterdefense system against bacterial hosts and might be used to improve human and animal health. For example, phages equipped with NAD^+^ synthetic pathway may be used for regulating gut microbiome or killing drug-resistant bacteria.

The host of many NAD-jumbo phages were Bacteroidota which are dominant symbiotic bacteria in the gut of animal and human, suggesting that NAD-jumbo phage may be an important component of intestinal microbiome. Moreover, NAD-jumbo phages have complemented NAD^+^ synthesis pathway with their bacterial hosts in animal and human gut, implying that NAD-jumbo phages may affect NAD^+^ synthesis of gut bacteria and even mammalian cells. Recent studies report a trans-kingdom cooperation of NAD^+^ synthesis between gut microbiome and mammalian cells wherein gut microbiota boost NAD^+^ biosynthesis of mammalian host, which suggesting the important role of this interaction in NAD-mediated antiaging or antitumor [42–44]. Therefore, the metabolic interaction between NAD-jumbo phages and gut core bacteria (such a Bacteroidota) discovered in the present study has some important implying for NAD-mediated human health. First, to improve the level of NAD^+^ in tissues for antiaging in human, future studies need to consider the effects of gut phage NAD^+^ metabolism on mammalian NAD^+^ biosynthesis. Second, NAMPT or pncA expression of gut bacteria promote mammalian NAD synthesis [45], which may lead tumor cell resistance and tumor cell survival or migration. Therefore, phage-encoded NAMPT or pncA genes likely affect microbe-mediated drug resistance in tumor microbiome [46].

Overall, there is still poor understanding on jumbo phages, and our expanded high-quality jumbo phages provide important resources for jumbo phage biology. Al-Shayeb et al. suggested that jumbo phages are completely different from small phages and more similar to symbiotic bacteria, blurring the distinctions between living and non-living [9]. Our findings reinforce the point given that one of the characteristics of life is the ability to producing metabolic energy such as NAD^+^ to sustain themselves. Given that jumbo phages have the potential to independently synthesize energy matter NAD^+^, we propose that NAD-jumbo phage are ancient and plays a role in the origin of life [47].

## Methods

### Metagenomic samples

A total of 424 cattle, 140 sheep, 169 pig, 29 deer, and 193 horse gut metagenomes, comprising 418 cattle, 66 sheep, 29 deer, 163 pig, and 116 horse metagenomes from the NCBI or ENA public databases, and 6 cattle, 6 pig, 74 sheep and 77 horse metagenomes from this study (Table S1). The 6 cattle metagenomic samples were from adult Holstein cattle, including 3 colonic contents and 3 rumen contents. The 6 pig metagenomic samples were from adult Duroc, including 3 colonic contents and 3 stomach contents. The 74 sheep metagenomic samples were from Kazakh sheep, including 33 rumen contents, 5 reticulum contents, 6 omasum contents, 6 abomasum contents, 6 ileal contents, 6 cecum contents, 6 colon contents and 6 rectal contents. All samples in this study were collected from slaughterhouse in Shihezi City, China. In addition, the 77 horse metagenomic samples were from the feces of thoroughbred horse. These samples were collected from the Hazanat International Racecourse in Mulei City, China. All samples were frozen in liquid nitrogen immediately after collection in sterile centrifuge tubes and stored at –80°C until genome extraction. All procedures involving animals were approved by the Animal Care Committee of Shihezi University. The study was performed in accordance with the ethical standards established in the 1964 Declaration of Helsinki and subsequent amendments.

### DNA extraction and sequencing

To obtain high-quality microbial DNA from the gut contents, the DNA of the samples was extracted by cetyltrimethyl ammonium bromide (CTAB) method [48]. The quality, integrity and potential contamination of extracted DNA were evaluated by agarose gel electrophoresis and pulsed field gel electrophoresis. Meanwhile, the Nanodrop Kit (Implen, CA, USA) and Qubit®2.0 fluorometer (Life Technologies, CA, USA) were used for purity determination and precise quantification of the extracted DNA. According to the manufacturer’s protocol, illumina sequencing libraries were prepared with a NEBNext®Ultra™DNA Library preparation kit (New England Biolabs, USA) of 500 mg DNA samples from each sample. These libraries were purified using the AMPure XP system (Beckman Coulter, Brea, CA, USA) and quantitatively analyzed using the Agilent 2100 bioanalyzer and real-time fluorescent quantitative PCR. All libraries were sequenced using Illumina NovaSeq 6000 platform with a reading length of 150 base pairs (PE150).

### Data quality control and phage genome identification

To identify phage sequences in cattle, sheep, horse, pig and deer gut samples, we performed a rigorous pipeline for phage genome assembly to avoid many false positives in the expected metagenome (Figure S1) [9, 49]. Obtaining high-quality metagenomic phage contigs is a critical step in subsequent analysis, we quality-controlled these data using Fastp software (Version: 0.20.1) and aligned them with cattle, sheep horse, pig and deer reference genomes using bowtie2 (Version: 2.2.4) to remove host sequences, respectively [50]. Next, the cleandata were assembled by the software metaSpades (Version: SPAdes-3.11.1) to obtain genomic contigs [51]. These genomic contigs were screened for metagenomic virus contigs using virFinder [52], VirSorter 2 [53], VIBRANT [54] and CheckV [55]. When using VirSorter, only contigs classified as 1, 2, 4 or 5 Category were considered. For VirFinder, we retained these contigs with scores > 0.9 and p < 0.01 and linear contigs >5 kbp in length and circular contigs > 1.5 kbp in length. These viral contigs were de-redundant using dRep software (Version: dRep v3.2.2, parameters: ––S_algorithm ANImf, ––S_ani 0.95, ––cov_thresh 0.8) [56]. Then only those sequences that VIBRANT identified as viruses with genomes size > 200 kbp in length were retained. After filtering, we obtained a total of 1001 jumbo phage genomes from the animal animal gut. Then, these phages were evaluated for quality using CheckV and the high quality and completed genomes (genome completeness > 90%) were retained for subsequent analysis. A total of 668 high quality jumbo phage genomes were obtained by using these stringent selecting criteria (Additional file 1: Fig. S1).

### Protein function annotation

Prodigal software using genetic code 11 (-m –g 11 –p single) (Version: V2.6.3) was used to predict ORFs encoded by 668 jumbo phage [57]. Further function annotation of protein sequences encoded by jumbo phage was performed by viral homolog (VOG) [58], KEGG [59] and Pfam [60]. tRNAscan-s.e.v.2.0 with bacterial model was used to identified tRNA sequences in jumbo phages and their hosts [61]. For detecting the presence of conserved tRNAs in jumbo phages, we used MEGA7.0 software to align the tRNA of jumbo phage with the corresponding tRNAs sequences from bacteria [62].

### Jumbo phage classification and phylogenetic analyses

To understand the taxonomic information of 668 jumbo phages, we downloaded the phage genome from the RefSeq prokaryotic database (Prokaryotic Viral RefSeq 211) and reported jumbo phages. The distance-based hierarchical clustering and virus classification prediction of protein-sharing networks of 668 jumbo phages and Reference phages were performed for population analysis using vConTACT2 (v0.11.3) [63]. The predicted protein sequences were subjected to all-verse-all BLASTP using DIAMOND (more sensitive mode, identity 25%, query cover 50%). Protein clusters were generated using MCL and viral clusters were generated using ClusterONE in vConTACT2 pipeline.

Based on functional annotations on the ORFs using eggnog-mapper, we select major capsid protein (MCP) and terminase large subunit (TerL) protein of jumbo phages in this study and previously reported [9]. All predicted MCP and TerL protein sequences were examined using PfamScan (v1.6) against Pfam 35 and manual check. DIAMOND BLASTP (-e 1e-10 ––more-sensitive) was used to search reference sequences against VOG 216 database. The host assignment of reference viruses was based on Virus-Host DB (https://www.genome.jp/virushostdb/) [64]. MCP and TerL protein sequences were aligned using linsi mode of the MAFFT (v7.310) software [65]. The alignments were trimmed using trimAL (v 1.4.rev15) to remove position with gaps over 90% sequences [66]. The Maximum-likehood phylogenetic trees of MCP and TerL protein sequences was computed by IQ-TREE (v2.0.3) after extended model selection (MCP LG+F+R9; TerL LG+F+R10), and support for nodes was evaluated with 1001 replicates for ultrafast bootstrap [67].

### Taxonomy of jumbo phage

The prokaryotic virus from NCBI GenBank, RefSeq and VOG database were constructed as a reference database using custom scripts. The ORFs encoded by jumbo phage were predicted by Prodigal software (Version: V2.6.3, parameter: –p meta) and constructed into a protein database using DIAMOND (Version: v0.9.9) [57]. The protein database was aligned with the reference database using DIAMOND BLASTp (Parameters: blastp –e 1e-5 ––query-cover 10 ––subject-cover 10) to obtain taxonomy of each ORFs encoded by jumbo phage [68]. We assigned taxonomic information of each jumbo phage based on the majority taxonomic assignment [49]. 668 Jumbo phage without the majority taxonomic assignments or with less than two assigned ORFs were considered as unassigned phages [49].

### Host prediction

We predicted CRISPR sequences in microbial metagenome-assembled genomes (MAGs) from the gut microbiome of animal (Supplementary dataset 1) to obtain CRISPR spacer sequences by using MinCED software (Version: V.0.4.2), a software based on CRT, to predict the possible host of huge phages [69]. The taxonomic assignment of the MAGs was generated by GTDB-tk (classify_wf workflow with r207 database, v2.1.0) and was converted into the corresponding NCBI phylum taxonomy [70]. In all, we extract 265818 CRISPR spacers within 12674 MAGs. CRISPR spacers sequences were queried with jumbo phage using blastn with E-value of <10−5, identity >95% and coverage >80% (with ≤1 mismatch over the whole spacer) [71]. The host of jumbo phage were assigned to CRISPR spacers taxonomy based on blast hits with the highest bit scores.

### Evolutionary analysis and protein structure prediction of NAMPT and NadD

The NAMPT and NadD sequences in NAD-jumbo phages were aligned using MUSCLE [72]. DNA alignments were further polished using trimAl [66]. The Maximum-likehood phylogenetic trees of NAMPT and NadD protein sequences was computed by IQ-TREE (v2.0.3) after extended model selection (MCP LG+F+R9; TerL LG+F+R10), and support for nodes was evaluated with 1001 replicates for ultrafast bootstrap. For RoseTTAFold predictions, run_e2e_ver.sh script was used to predict the three-dimensional structure of NAMPT and NadD proteins [73]. The three-dimensional structure of NAMPT and NadD encoded in NAD-jumbo phages were compared with those encoded in bacteria [74, 75] using Foldseek’s easy-to-search command default parameter [76].

## Data availability

All the original sequences obtained and published data in this work have been uploaded to public databases, and detailed project numbers can be found in supplementary table S1. All the data can also be obtained from the corresponding author upon reasonable request or from the included supplementary information files.

## Contributions

C.L., K.L., W.N., M.L. and S.H. designed the research. C.L., K.L. and C.G. performed analysis. C.L., K.L., M.L., P.Z., Z.S., X.L. and L.W. performed the collection of samples. W.N., M.L. and S.H. supervised the study. C.L., K.L., W.N., M.L. and S.H. wrote the manuscript. C.L., M.L. L.C and S.H. improved the manuscript. All authors approved the final version of the manuscript.

## Supplementary Figures 1-9

Figure S1. Pipeline for high-quality jumbo phage construction.

Figure S2. Genome size distribution of jumbo phage in the gut of animal.

Figure S3. Distribution of genome completeness and classification of jumbo phage genomes into quality tiers. The abscissa represents the genomes length, and the ordinate represents the genomes completeness.

Figure S4. Genome circos map of 716.8kbp and 713.6kbp jumbo phages.

Figure S5. Sequence alignment of tRNAs encoded by jumbo phages and hosts.

Figure S6. Network images of the newly assembled jumbo phages based on shared protein ortholog families. vConTACT2 v0.11.3 was used for clustering each phage. Our jumbo phages were highlighted in different colors.

Figure S7. Genetic maps of NAD-jumbo phage. The white graph represents the unannotated protein.

Figure S8. The phylogeny of ligA/B gene encoded by jumbo phage. The ligA/B gene encoded by the jumbo phage in this study are shown in green.

Figure S9. NAD^+^ biosynthesis pathway are complementary between NAD-jumbo phages and their bacterial hosts. NAD-jumbo phages from human gut are paired with their hosts. The purple circles represent the presence of the corresponding enzyme in the NAD-jumbo phage. The green circles represent the presence of the corresponding enzyme in the host. The space circle represent the absence of the corresponding enzyme in the genome.

## Supplementary Tables 1-12

Supplementary Table 1: All detailed information on samples and metagenomic data.

Supplementary Table 2: Basic characteristics of the assembled jumbo phages.

Supplementary Table 3: Putative transfer RNA information encoded by the jumbo phages.

Supplementary Table 4: Functional annotation of our assembled jumbo phage genomes.

Supplementary Table 5: Clade distribution of our assembled jumbo phages in phylogenetic trees.

Supplementary Table 6: Proto-spacer and host predicted in the jumbo phage genomes.

Supplementary Table 7: Basic information and gene function annotation of NAD-jumbo phages.

Supplementary Table 8: The NAD-jumbo phages encode a large number of NAD-depended enzymes and NAD^+^ cofactors.

Supplementary Table 9: Jumbo phages and NAD-jumbo phages assembled from the human body samples.

Supplementary Table 10: Basic information of jumbo phages and NAD-jumbo phages assembled from metagenomic data of different environments.

Supplementary Table 11: Genomic annotation of NAD-jumbo phages from different environmental metagenomic samples.

Supplementary Table 12: Host prediction of NAD-jumbo phages assembled from human metagenomic samples.

Supplementary dataset 1: animal gut MAG dataset contains 29, 599 genomes of bacteria and archaea.

Supplementary dataset 2: JumbophageDB includes 1864 draft genomes of jumbo phages.

