## Supplementary material for "Jumbo phages possess independent synthesis and utilization systems of NAD^+^": Figure S1-S9: Figure S1-S9.pdf

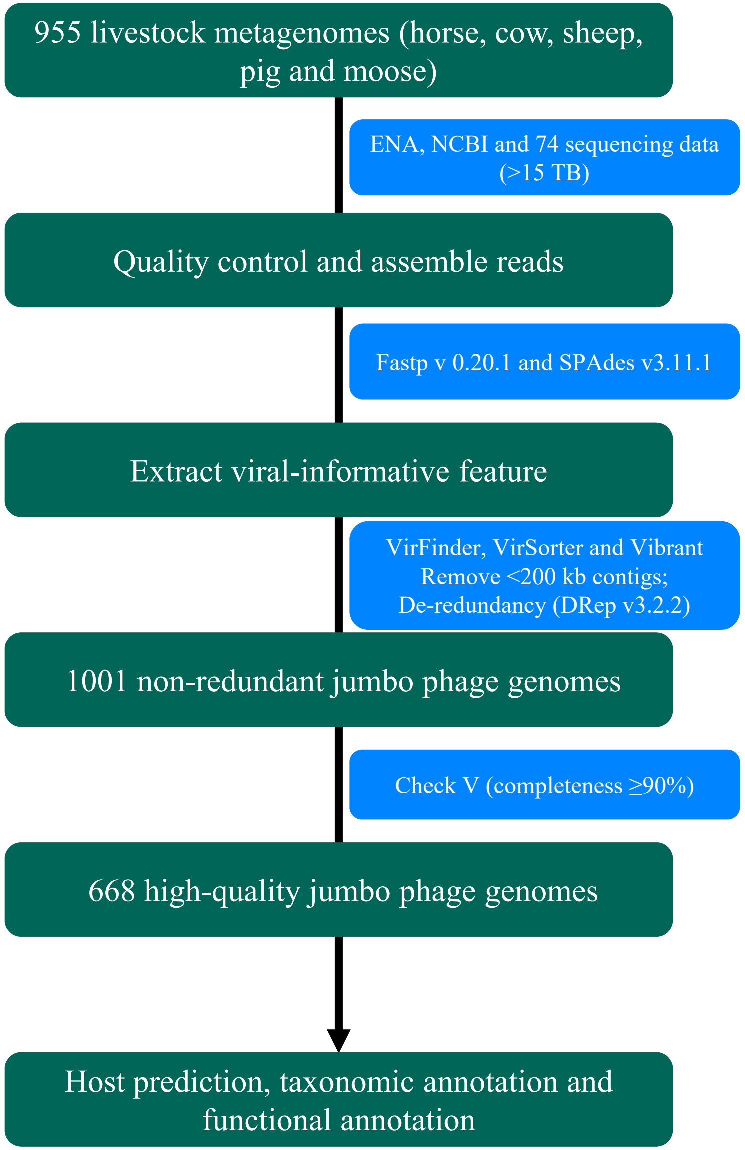


**Figure S1. Pipeline for high-quality jumbo phage construction.**

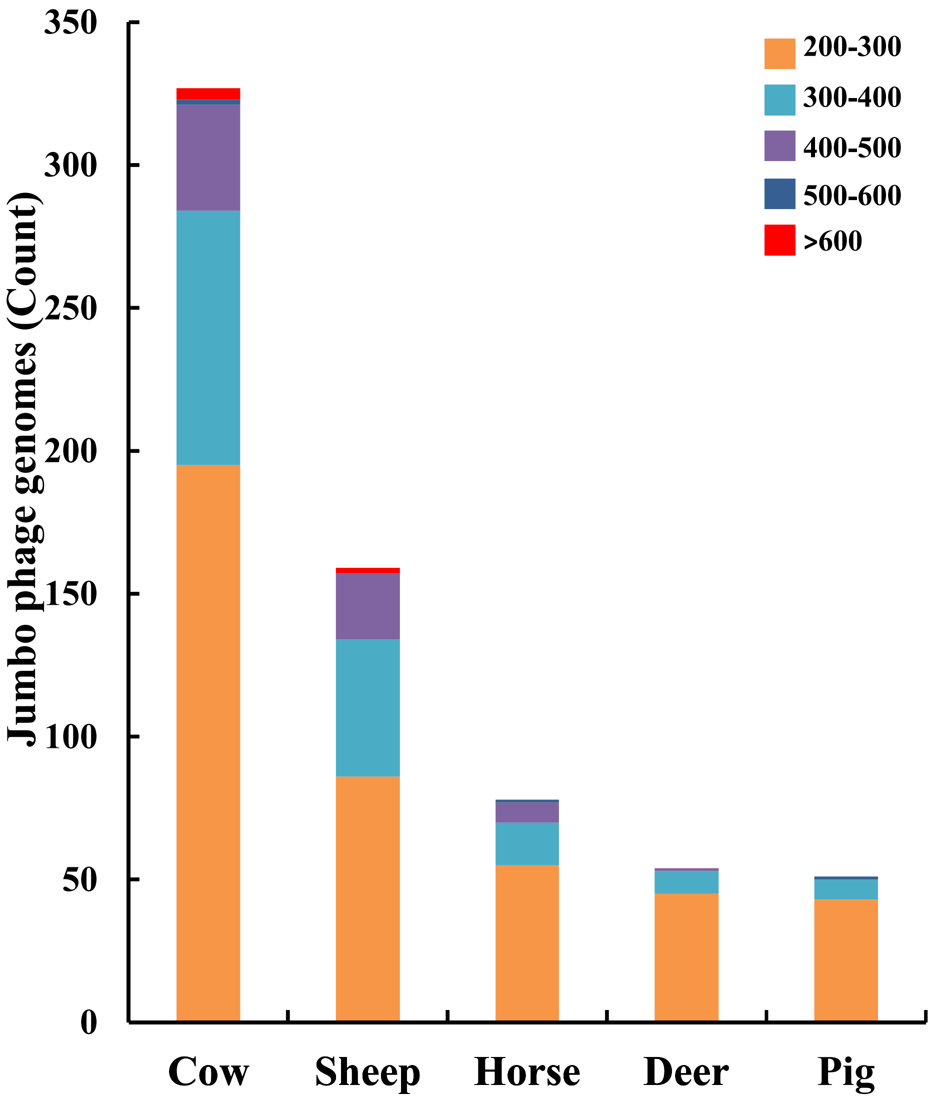


**Figure S2. Genome size distribution of jumbo phage in the gut of livestock.**

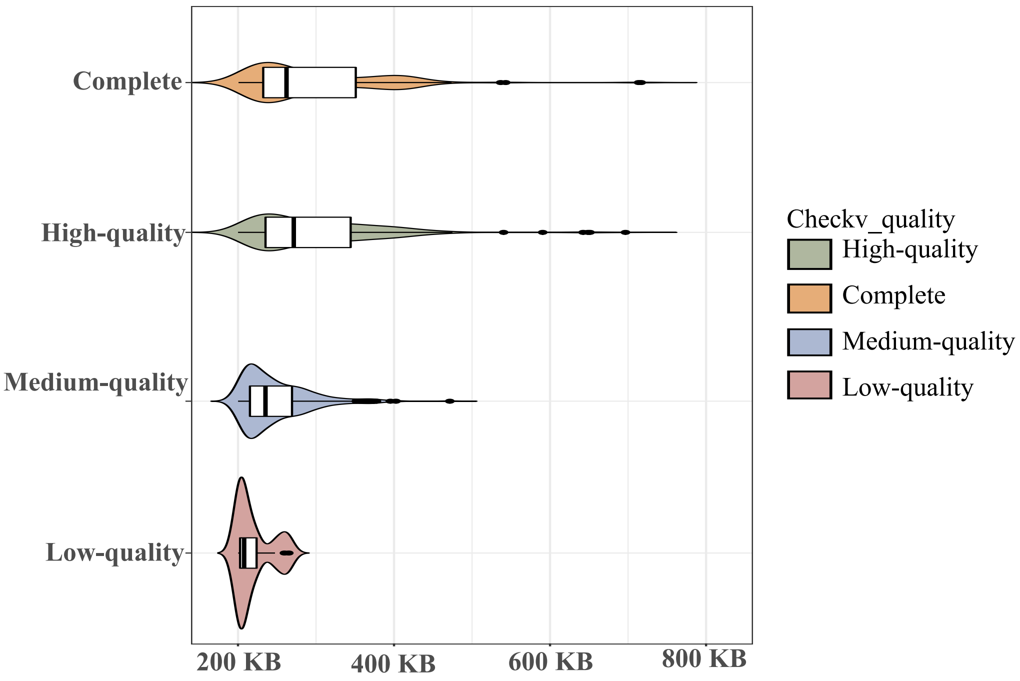


**Figure S3. Distribution of genome completeness and classification of jumbo phage genomes into quality tiers.** The abscissa represents the genomes length, and the ordinate represents the genomes completeness.

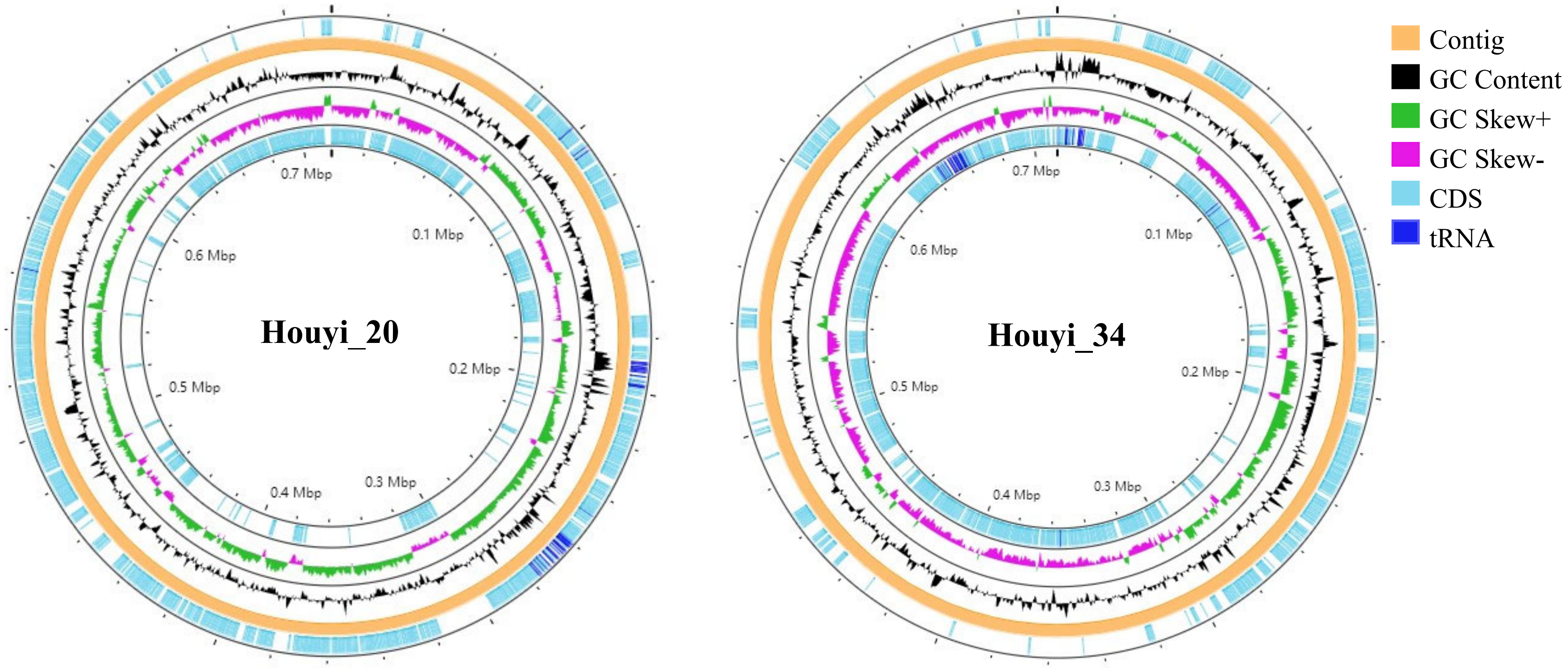


**Figure S4. Genome circos map of 716.8kb and 713.6kb jumbo phage.**

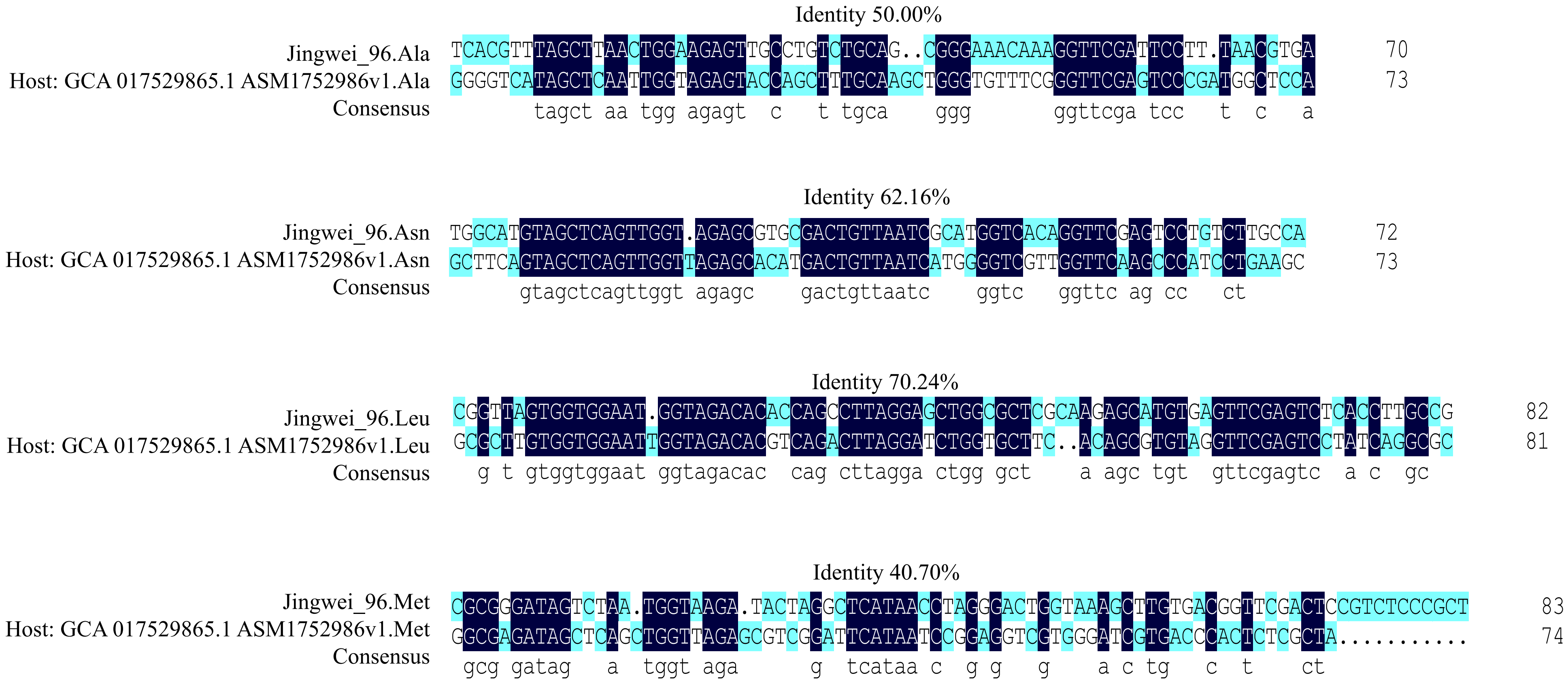

**Figure S5. Sequence alignment of tRNAs encoded by jumbo phage and host.**

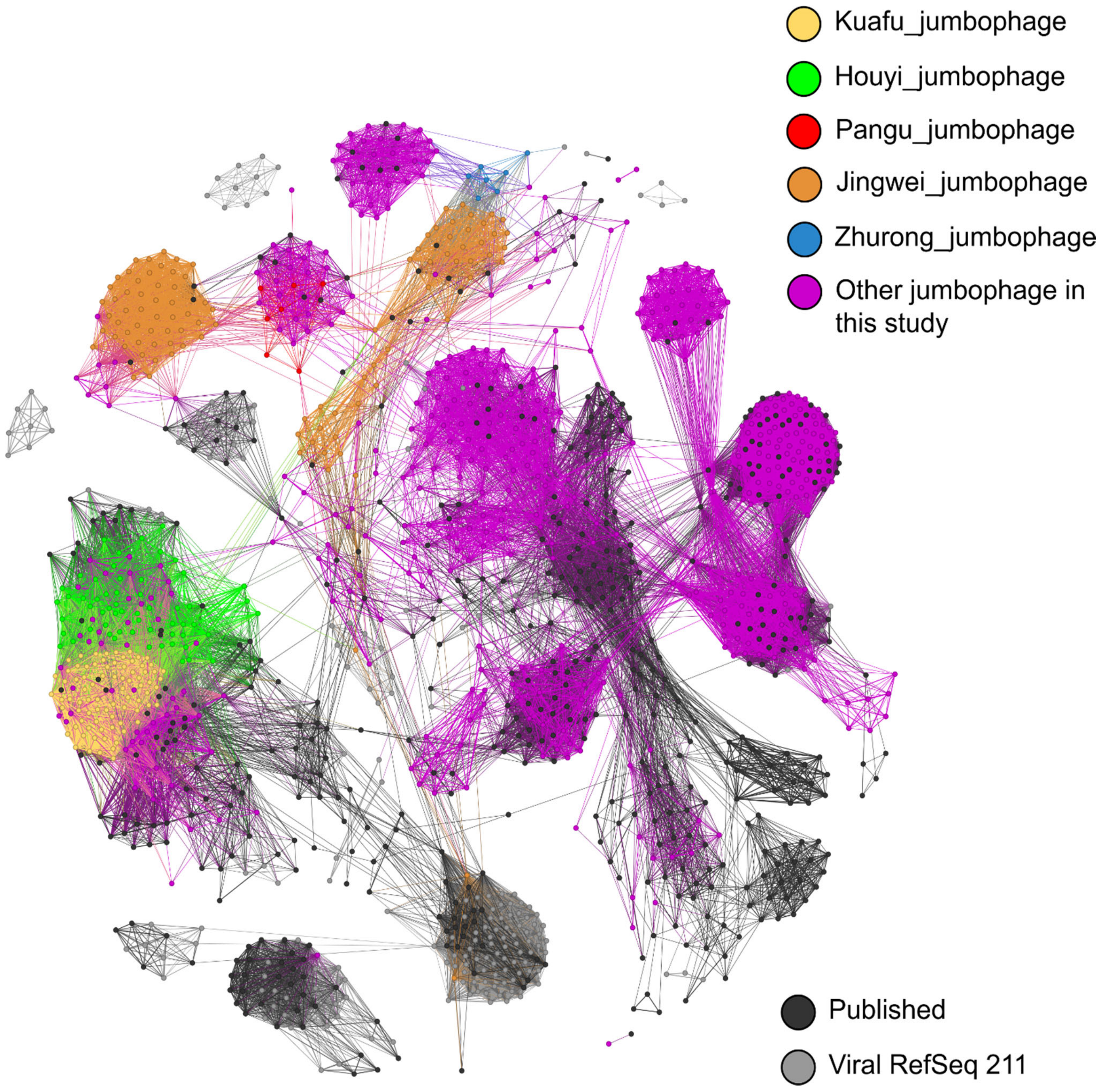


**Figure S6. Network images of the newly assembled jumbo phages based on shared protein ortholog families.** vConTACT2 v0.11.3 was used for clustering each phage. Our jumbo phages were highlighted in different colors.

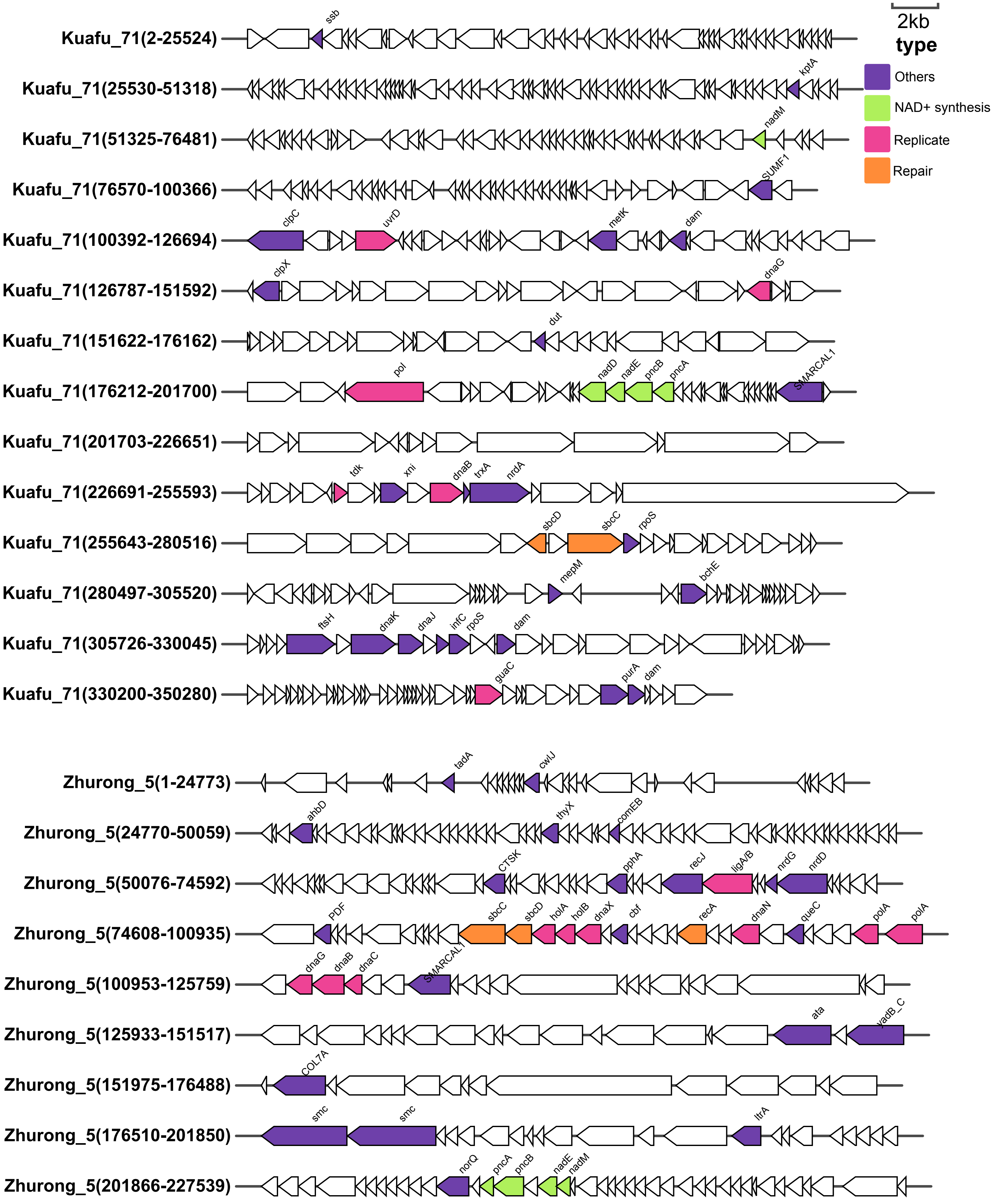


**Figure S7. Genetic maps of NAD-jumbo phage.** The white graph represents the unannotated protein.

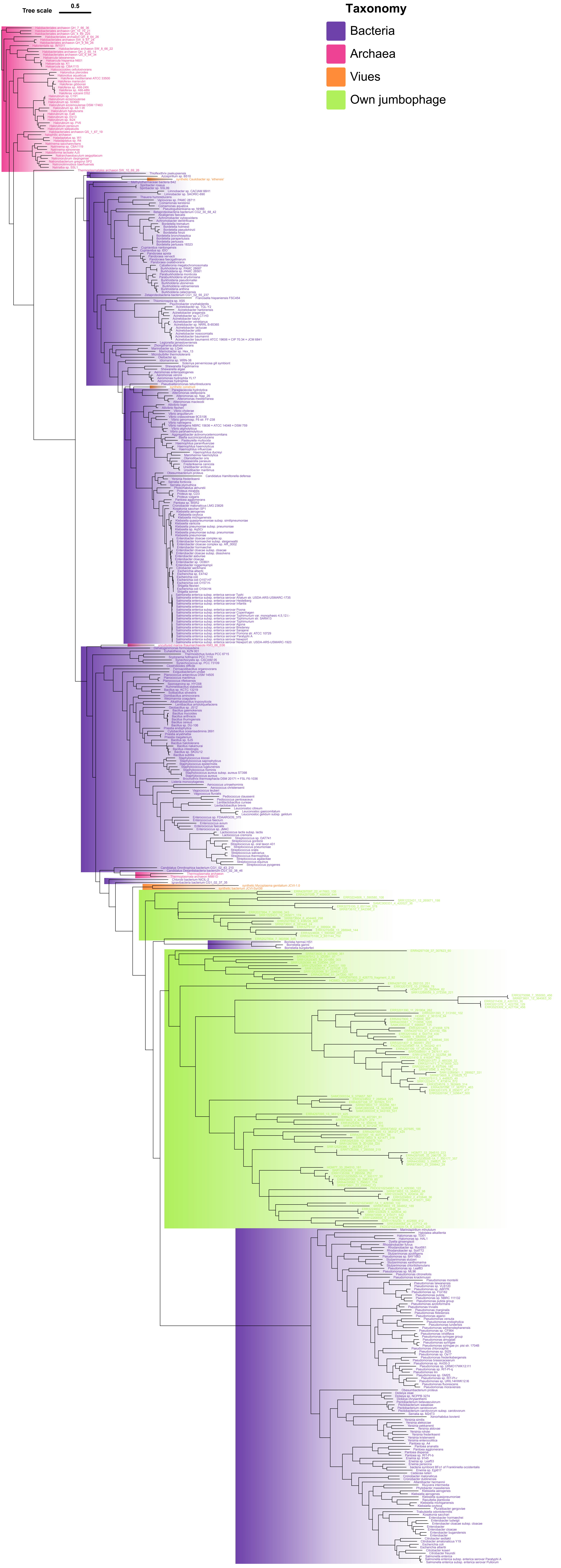


**Figure S8. The phylogeny of ligA/B gene encoded by jumbo phage.** The ligA/B gene encoded by the jumbophage reported here are shown in green.

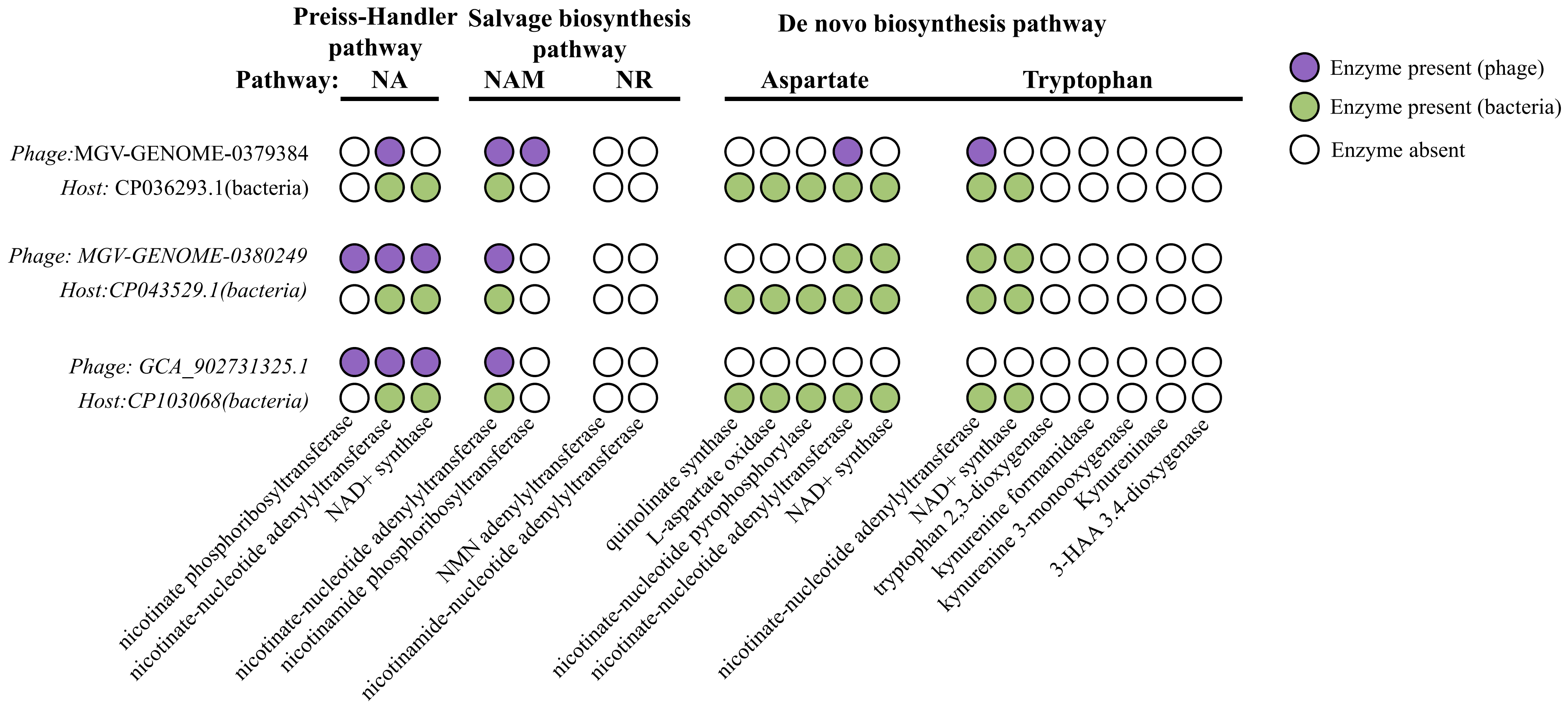


**Figure S9. The human body NAD-jumbo phage complements its host in the NAD synthesis pathway.** The white circles represent the absence of the corresponding enzyme in the genome, the green circles represent the presence of the corresponding enzyme in the host, and the purple circles represent the presence of the corresponding enzyme in the NAD-jumbo phage.
